# Motor learning in real-world pool billiards

**DOI:** 10.1101/612218

**Authors:** Shlomi Haar, Camille M. van Assel, A. Aldo Faisal

**Author notes:** **Corresponding authors:** (AAF) and (SH). **Declaration of Interests**: The authors declare no competing financial interests. **Contributions:** SH and AAF conceived and designed the study; SH, CVA, and AAF developed the experimental setup; SH and CVA acquired the data; SH and AAF analysed and interpreted the data; SH drafted the paper; SH and AAF revised the paper.

## Abstract

The neurobehavioral mechanisms of human motor-control and learning evolved in free behaving, real-life settings, yet this is studied mostly in reductionistic lab-based experiments. Here we take a step towards a more real-world motor neuroscience using wearables for naturalistic full-body motion-tracking and the sports of pool billiards to frame a real-world skill learning experiment. First, we asked if well-known features of motor learning in lab-based experiments generalize to a real-world task. We found similarities in many features such as multiple learning rates, and the relationship between task-related variability and motor learning. Our data-driven approach reveals the structure and complexity of movement, variability, and motor-learning, enabling an in-depth understanding of the structure of motor learning in three ways: First, while expecting most of the movement learning is done by the cue-wielding arm, we find that motor-learning affects the whole body, changing motor-control from head to toe. Second, during learning, all subjects decreased their movement variability and their variability in the outcome. Subjects who were initially more variable were also more variable after learning. Lastly, when screening the link across subjects between initial variability in individual joints and learning, we found that only the initial variability in the right forearm supination shows a significant correlation to the subjects’ learning rates. This is in-line with the relationship between learning and variability: while learning leads to an overall reduction in movement variability, only initial variability in specific task-relevant dimensions can facilitate faster learning.

## Introduction

Motor learning is a key feature of our development and daily lives, from a baby learning to crawl, an adult learning a new sport, or a patient undergoing rehabilitation after a stroke. The process of learning a real-world motor skill is usually long and complex, and difficult to quantify as tasks are naturally unconstrained and highly variable. Most of the motor learning literature focuses on relatively simple tasks, performed in a laboratory setup or even within an MRI scanner, such as force-field adaptations^1–4^, visuomotor perturbations^5–9^, and sequence-learning of finger tapping or pinching tasks^10–13^. These laboratory-based learning tasks enable us to isolate specific features of motor learning and dissect them individually, thus provide elegant experiment designs to verify the experimenters’ hypothesis. As a result, motor learning in the real world is rarely studied. While laboratory-tasks play an important role in our understanding of sensorimotor control and learning, they address a very restricted range of behaviours that do not capture the full complexity of real-world motor control and may overlook fundamental principles of motor control and learning in real-life^14,15^.

Neurobehavioral mechanisms are subject to evolutionary selection pressures and survive only if they are relevant in natural tasks. Thus, studying operation in natural contexts allows us to evaluate mechanisms the nervous system has been designed for^16,17^. E.g. in sensory neuroscience the use of natural sensory stimuli has led to a revolution of our mechanistic understanding of perception^18,19^. Over the past decade, there were a few notable efforts to study motor learning in unconstrained tasks. One line of research devised more complex tasks for skill learning^20–22^ (e.g. skittles) which were implemented as computer-based gamified tasks that emulate real-world tasks. Others moved away from the computer screen but were still highly constrained; e.g. throwing a frisbee while the subject’s trunk is strapped to the chair to prevent trunk movement^23^. Another line of inquiry used free-behaviour in real-world tasks such as tool-making or juggling^17,24–27^. In these studies, we and others focused on anatomical and functional MRI measurable changes following learning. In a 3-year long complex tool-making apprenticeship experiment^17^ we were able to quantify changes in motor control precision and improvements of task outcomes, but given the 100s of hours of training involved and complexity of the task itself, we were not able to record trial-by-trial learning effects. Therefore, the findings and insights of learning studied in the computational motor control literature – at the level of actual changes in motor coordination and control policies – has received little attention in previous real-world learning studies.

One of the main challenges of understanding real-world behaviour and specifically motor learning is to identify underlying simplicities in a highly variable stream of movements that are not well constrained. We take a data-driven approach to analyse a real-world task and thereby illustrate a process by which one can investigate a real-world motor learning task in a principled manner. We aim to inject as few assumptions a priori as possible about task-relevant joints or mechanisms of learning, instead we aim to use plausible methods to reveal these to us. The paradigm we choose is the game of pool table billiards. Pool is a real-world task that involves many different sub-tasks (precision, alignment, ballistic movements, high-level sequential planning and sequencing of shots and ball positions) which requires advanced skills – hence it is a highly competitive sport. While our ultimate goal is to move to a data analysis framework for real-world motor learning in arbitrary tasks, we find that the game of pool offers a useful intermediate goal that frames the behavioural data in a way that makes our approach amenable to be understood in the currently dominant framework of laboratory-tasks and measures. Pool billiards has natural spatial constraints (the area in and around the pool table), divisibility of behaviour into trials (shots of the ball) allowing us to visualize results in the same framework of lab-based motor learning tasks and a clear outcome (sink the ball into the pocket). Subjects had to do a pool shot to put the ball in the pocket using unconstrained full-body, self-paced movement, with as many preparatory movements as the subject needs for each shoot, the only constraints arose from the placement of the white cue ball which the subjects shoot with the cue stick and the red target ball (that needs to go into the pocket). We implemented this as a real-world experiment, effectively only adding sensors to the subject and the pool table, i.e. subjects use the normal pool cue, balls, and pool table they would in a leisure setting and thus carry out natural motor commands, receive the natural somatosensory feedback and experience the same satisfaction (rewards) when they put the ball in the pocket as this is a real-world task. Crucially, the skill of subjects in putting the ball into the pocket is learnable in the time course of 1-2 hours, allowing us to record and analyse the experiments as one session.

To tackle the complexity of the high dimensional full-body motor control and task-space (game objects) movement, we recorded continuously the full-body movement during the entire learning period (about an hour and a half) and measured balls movements automatically on the table. EEG activity was also recorded during the task via mobile brain imaging, but to focus here on the motor kinematics learning we chose to report the neural activity results elsewhere^28^. We quantify the trends in full-body movement and task performance separately during the entire learning process, and look for correlations between the changes in the body movement and the performance in the task.

We structured the results as follows: We ground our results in previous work on laboratory-tasks, to show that our unconstrained task and its task goal (directional error of the target ball relative to the pocket it is meant to go in) displays the well-known features of human motor learning, namely learning curves with characteristic double exponential shape. We then characterize full-body movement structure during the task, and how learning changes the kinematics of every of the measured 18 joints. In our analysis, we will alternate between taking a data-driven view that attempts to be task-ignorant to identify underlying simplicities indicative of biological mechanisms in the data, and a task-based view to interpret the data-driven findings by using task-domain knowledge. Finally, we compare across subjects to characterize how their performance, motor variability, and learning rates are linked.

## Results

30 right-handed volunteers, with little to no previous experience playing billiards, performed 300 repeated trials (6 sets of 50 trials each with short breaks in-between) where the cue ball and target ball were placed in the same locations, and subjects were asked to shoot the target ball towards the far-left corner pocket (Figure 1A). During the entire learning process, we recorded the subjects’ full-body movements with a ‘suit’ of inertial measurement units (IMUs; Figure 1B), and the balls on the pool table were tracked with a high-speed camera to assess the outcome of each trial (Figure 1C).

**Figure 1.**
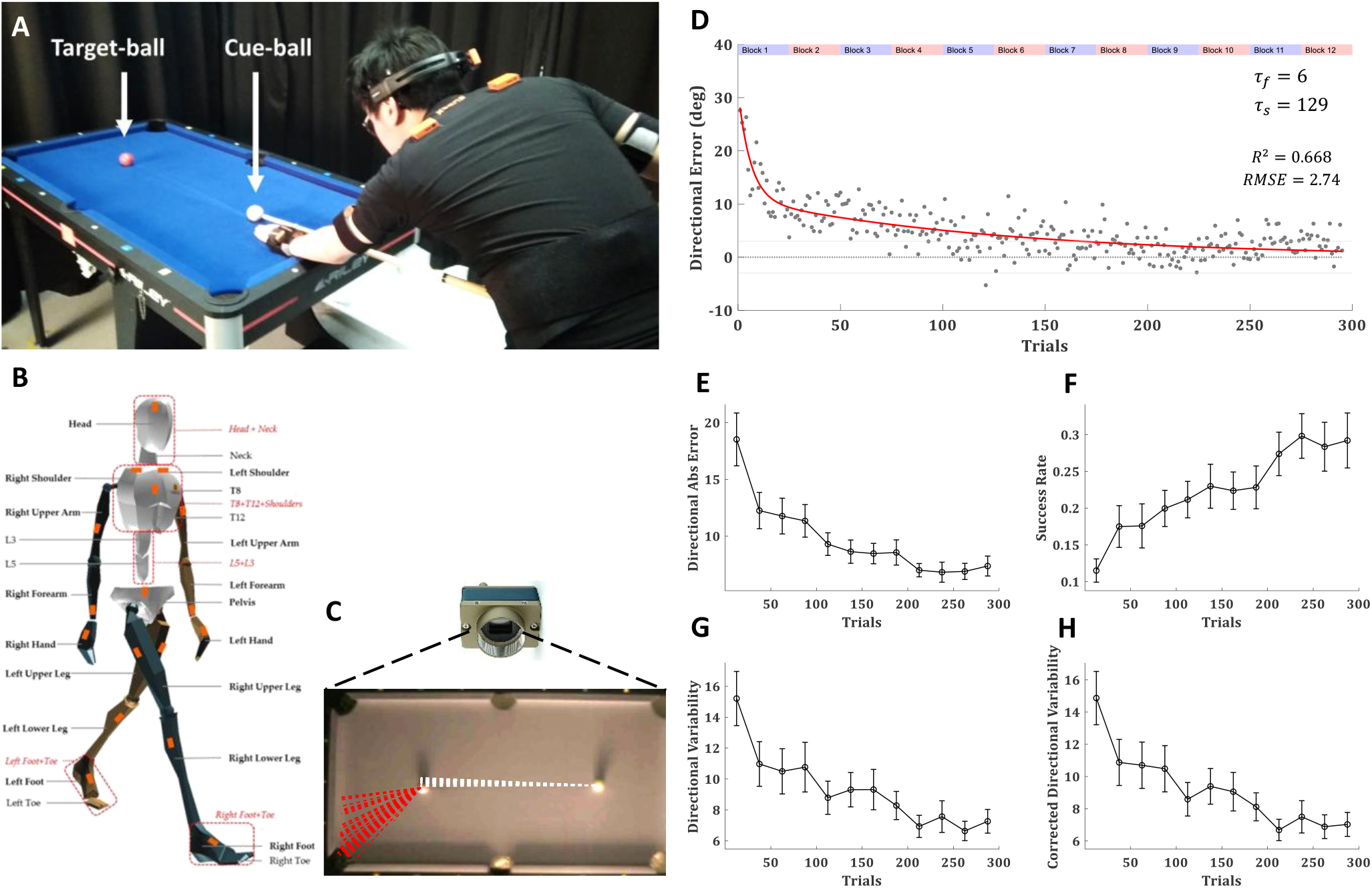
*Experimental setup and task performance*. (**A**) 30 right-handed healthy subjects performed 300 repeated trials of billiards shoots of the target (red) ball towards the far-left corner. (**B**) Full body movement was recorded with a ‘suit’ of 17 wireless IMUs (Xsens MVN Awinda). (**C**) The pool balls were tracked with a high-speed camera. Dashed lines show the trajectories of the cue (white) and target (red) balls over 50 trials of an example subject. (**D**) The trial-by-trial directional error of the target-ball (relative to the direction from its origin to the centre of the target pocket), averaged across all subjects, with a double-exponential fit (red curve). The time constant of the fast and slow components were 6 and 129 trials, respectively. Grey lines mark the range of successful trials (less than 3 degrees form the centre of the pocket). (**E**) The mean absolute directional error of the target-ball. (**F**) The success rates. (**G**) directional variability. and (**H**) directional variability corrected for learning (see text). (**E-H**) presented over blocks of 25 trials, averaged across all subjects, error bars represent SEM.

### Movement and Learning in a real-world pool task

The ball tracking data showed learning curve for the decay in the directional error of the target ball (relative to the direction from its origin to the centre of the target pocket) over trials (Figure 1D). This learning curve was best fit with a double exponential curve (Supplementary Figure 1). The direction of the error in the initial trials was consistent across subjects as they tended to hit the centre of the target ball and shot it forward towards the centre of the table. For measuring success rates and intertrial variability we divided the trials into blocks of 25 trials (each experimental set of 50 trials was divided into two blocks to increase the resolution in time). To improve robustness and account for outliers, we fitted the errors in each block with a t-distribution and used the location and scale parameters (μ and σ) as the blocks’ centre and variability measures. The learning curve over blocks (Figure 1E) emphasised the reduction in the inter-subject variability during learning (decreasing error bars). The success rate over blocks (percentage of successful trials in each block; Figure 1F) showed similar learning to the directional error.

Learning was also evident in the intertrial variability in the shooting direction which decayed over learning (Figure 1G). Since learning also occurred within a block (especially during the first block) and the variability might be driven by the learning gradient, we corrected for it by calculating intertrial variability over the residuals from a regression line fitted to the ball direction in each block (while the learning curve is exponential, within the small blocks of 25 trials it is almost linear). This corrected intertrial variability showed only minor reduction in the initial blocks, relative to the uncorrected variability, and showed the same decay pattern over the learning (Figure 1H). Overall, the task performance data suggested that subjects reached peak performance by the fifth experimental set (blocks 9-10, trials 200-250) and were doing the same (or even slightly worse) on the last experimental set (blocks 11-12, trials 250-300). Thus, we refer to the last two experimental sets (blocks 9-12, trials 201-300) as the ‘learning plateau’, while being mindful that professional pool players train over months and years to improve or maintain their skills.

Kinematic data were recorded using a wearable motion tracking ‘suit’ of wireless IMUs, where individual wireless sensors (matchbox-sized) were attached via Velcro to elastic straps fixed around the subjects’ body without constraining movement. The full-body kinematics were analysed in terms of joint angles using 3 degrees of freedom for each joint following the International Society of Biomechanics (ISB) recommendations for Euler angle extractions of Z (flexion/extension), X (abduction/adduction), and Y (internal/external rotation). Note, this standard approach includes hinge joints of the body which have only 1 degree of freedom being recorded as 3 Euler angles. The full-body movements were analysed over the angular joint velocity profiles of all joints. The data allowed us to reconstruct the full-body pose at any given moment, which we checked for visual correctness on a subject-by-subject basis against video ground truth. However, we chose not to look at joint angle’s probability distributions, as those are more sensitive to potential drifts in the IMUs (and contain small changes not spottable by the human eye). We previously showed that joint angular velocity probability distributions are more subject invariant than joint angle distributions suggesting these are the reproducible features across subjects in natural behavior^29^. In the current study, this robustness is quite intuitive: all subjects stood in front of the same pool table and used the same cue stick, thus the subjects’ body size influenced their joint angles distributions (taller subjects with longer arms had to bend more towards the table and flex their elbow less than shorter subjects with shorter limbs) but not joints angular velocity probability distributions (Figure 2).

**Figure 2.**
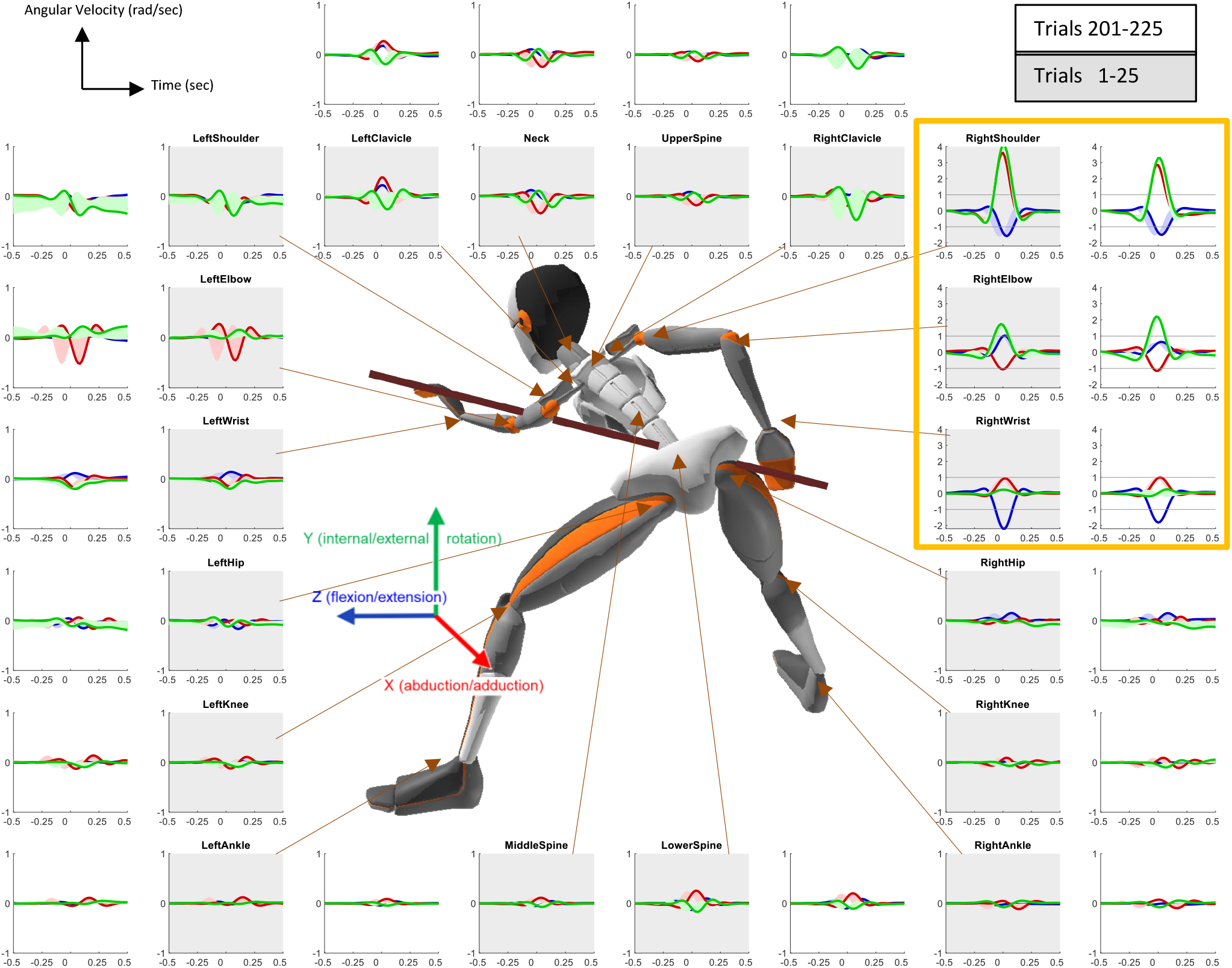
Angular velocity profiles. Angular velocity profiles in 3 degrees of freedom (DoF) for each joint (blue: flexion/extension, red: abduction/adduction; green: internal/external rotation) averaged across subjects and trials over the first block of trials (1-25) in the inner circle (grey background) and the first block after learning plateau (201-225) in the outer circle (white background). Shaded areas represent the standard error of the mean. Data was recorded in 60 Hz, X axis is in seconds, covering a 1 second window around the timepoint the cue hit the ball. Y axis is in radians per second. The joints of the right arm which do most of movement in the task are highlighted in orange box and have a different scale on the Y axis and grey line to indicate the Y axis limits of all other joints.

In the first step of our data-driven analysis, we wanted to identify the key joints for the task: we analysed the angular velocity profiles of all joints, averaged across the initial block trials of all subjects, and found that most of the movement is done by the right arm, and specifically in the right shoulder (Figure 2 inner circle). This is expected as all subjects were right-handed and used their right hand to hold the cue stick and make the shot. Taking domain understanding of the task into account, we can explain the shoulder movement by the naivety of the subjects, as pool billiards guidebooks^30–33^ emphasize that the shooting movement should be from the elbow down while the shoulder should be kept still. Correspondingly, the angular velocity profiles averaged across the initial block of the learning plateau (trials 201-225) showed similar distributions with an overall decrease in peak velocities relative to the initial trials but an increase in the peak angular velocity of ‘right elbow rotation’, which is the rotation between the upper arm and the forearm sensor and is equivalent to forearm supination (Figure 2 outer circle).

The angular velocities of all other joints were much smaller than those of the shooting (right) arm. For visibility, we increased the y-axis range of the right arm joints by a factor of 3 relative to all other joints (Figure 2). The high variance (relative to the mean) in some of the non-shooting arm joints that do move (such as the left elbow) suggests variability across trials and subjects in the movement of this joint and specifically in its timing relative to the shot. While some subjects in some shots had a small left elbow movement just before the peak of the shot, others had it shortly after. The sensor noise was much smaller than variability across trials and subjects, as demonstrated in a recent work where we provide the noise floor for the IMU, specifically, angular velocity precision evaluated against ground-truth marker-based optical motion tracking^34^.

To quantify the overall change in the within-trial variability structure of the body over trials, we use the generalised variance, which is the determinant of the covariance matrix^35^ and is intuitively related to the multidimensional scatter of data points around their mean. We measured the generalised variance over the angular velocity profiles of all joints and found that it increased rapidly over the first ~40 trials and later decreased slowly (Figure 3A). To understand what drives the generalised variance peak we plotted the variance-covariance matrixes of the first block, the second block (over the peak generalised variance), and ninth block (after learning plateaus) (Figure 3B). It shows that the changes in the generalised variance were driven by an increase in the variance of all DoFs of the right shoulder and the negative covariance between the abduction/adduction and internal/external rotation of the right shoulder to the flexion/extension of the right shoulder and wrist. The internal/external rotation of the right elbow showed a continuous increase in its variance, which did not follow the trend of the generalised variance.

**Figure 3.**
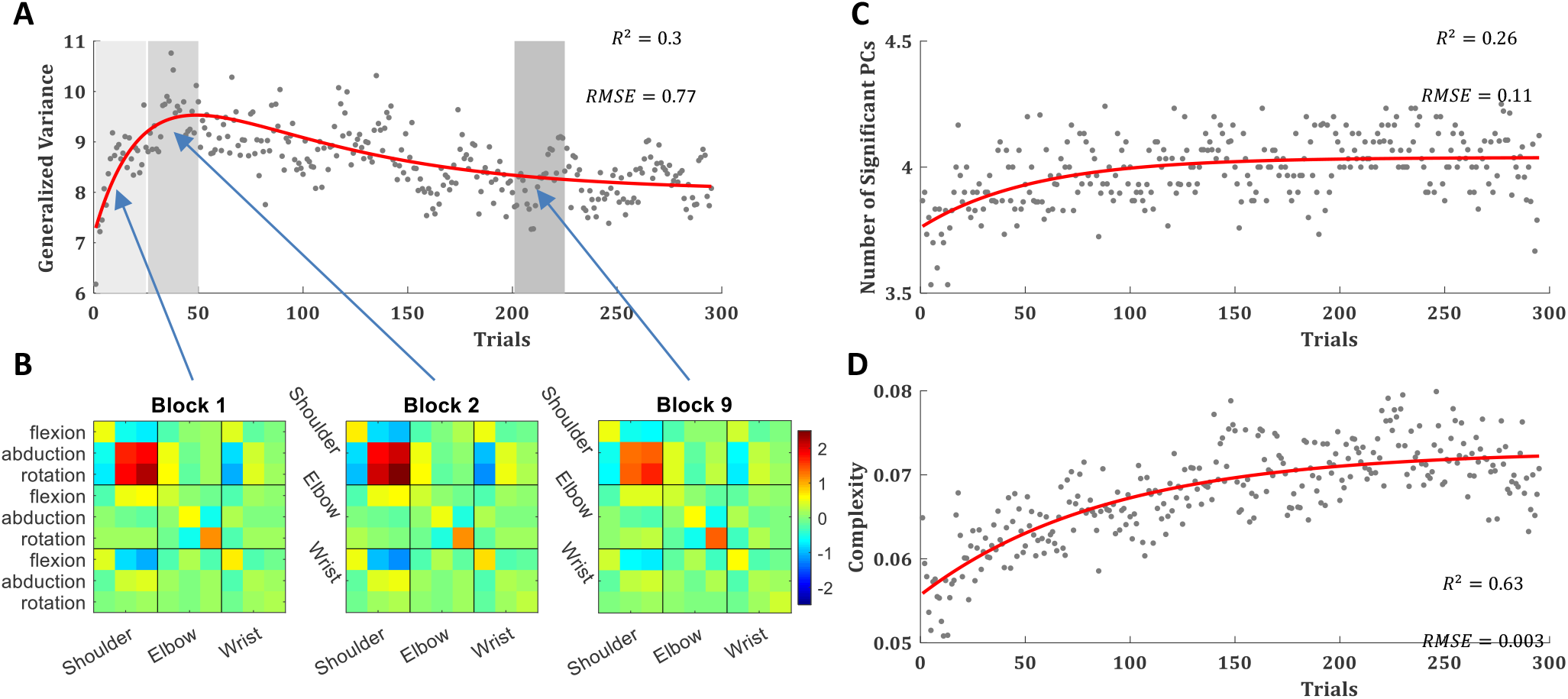
Variance and Complexity. (**A**) The trial-by-trial generalised variance, with a double-exponential fit (red curve). (**B**) The variance covariance matrix of the right arm joints angular velocity profiles averaged across subjects and trials over the initial block (trials 1-25), the second block (trials 26-50), in which the generalised variance peaks, and first block after learning plateau (block 9, trials 201-225). The order of the DoF for each joint is: flexion/extension, abduction/adduction, internal/external rotation. (**C**) The number of principal components (PCs) that explain more than 1% of the variance in the angular velocity profiles of all joints in a single trial, with an exponential fit (red curve). (**D**) The manipulative complexity (Belić and Faisal, 2015), with an exponential fit (red curve). (**A, C, D**) Averaged across all subjects over all trials.

Next we set to study the complexity of the movement – as defined by the number of degrees of freedom used by the subject – since the use of multiple degrees of freedom in the movement is a hallmark of skill learning^36^. For that purpose, we applied Principal component analysis (PCA) across joints for the angular velocity profiles per trial for each subject and used the number of PCs that explain more than 1% of the variance to quantify the degrees of freedom in each trial movement. While in all trials of all subjects most of the variance can be explained by the first PC (Supplementary Figure 2), there is a slow but consistent rise in the number of PCs that explain more than 1% of the variance in the joint angular velocity profiles (Figure 3C). The manipulative complexity, suggested by Belić and Faisal^37^ as a way to quantify complexity for a given number of PCs on a fixed scale (C = 1 implies that all PCs contribute equally, and C = 0 if one PC explains all data variability), showed cleaner trajectory with the same trend (Figure 3D). This suggests that over trials subjects use more degrees of freedom in their movement.

In the next step of our data-driven analysis, we wanted to identify signatures of learning joint-by-joint. For that, we defined a measure of task performance in single-joint space, which we named the Velocity Profile Error (VPE). VPE is the minimal correlation distances between the angular velocity profile of each joint in each trial to the angular velocity profiles of that joint in all successful trials (for more see methods). For all joints, VPE showed a clear pattern of decay over trials in an exponential learning curve (Figure 4A). We fitted it with a single exponential learning curve (see fits time constants and goodness of fit in Supplementary Table 1). A proximal-to-distal gradient in the time constant of these learning curves was observed across the right arm, from the shoulder to the elbow and the wrist rotation (Supplementary Figure 3). Intertrial variability in joint movement was measured over the VPEs in each block. Learning was also evident in the decay of the VPE intertrial variability during the learning over most joints across the body (Figure 4B).

**Figure 4.**
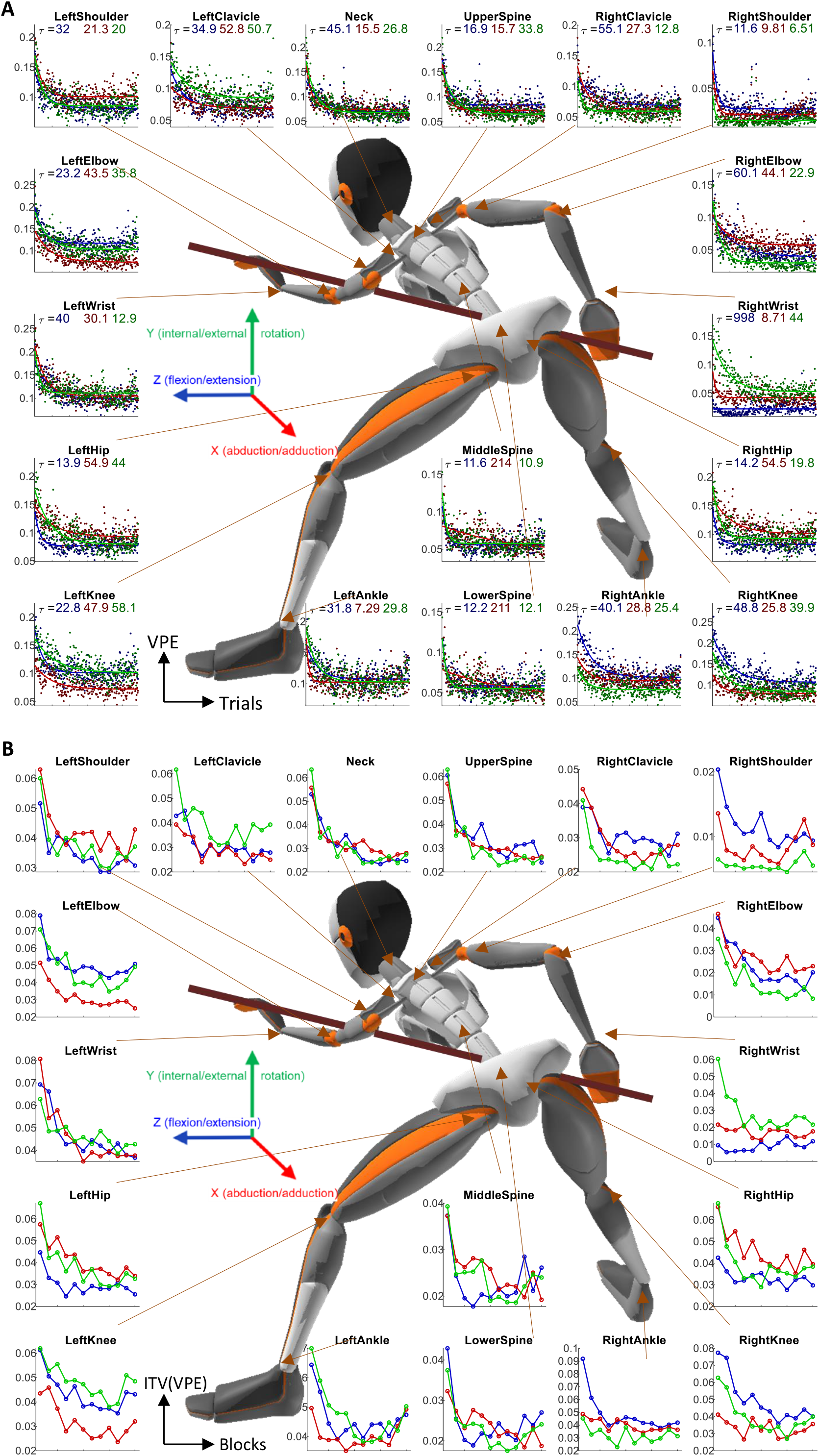
Learning over Joints. (**A**) Trial-by-trial Velocity Profile Error (VPE) for all 3 DoF of all joints, averaged across all subjects, with an exponential fit. The time constants of the fits are reported under the title. (**B**) VPE intertrial variability (ITV) over blocks of 25 trials, averaged across all subjects. The color code of the DoF is the same as in figure 2 (blue: flexion/extension; red: abduction/adduction; green: internal/external rotation).

### Inter-subject differences in variability and learning

In the final step of our data-driven analysis, we addressed the individuality of the subjects and looked across subjects for correlations between their task performance, learning rate, and joint movements. Since this is an exploratory study, all statistical tests are reported with caution and thus are not presented in the text but only in the figure (Figure 5) – where the readers can see the data points and make their judgment as for the true significance. The statistics are presented in Spearman rank correlation, to deal account for outliers and non-linear trends, and p-values are FDR corrected for multiple comparisons. The regression lines are presented only for visual account and include their 95% confidence intervals to address outlier biases. We found substantial differences between subjects in their initial errors, final errors, intertrial variability, and learning, which are overlooked in the group average results. One subject, who had low initial errors, showed no learning, i.e. did not reduce her error over trials from the first block (trials 1-25) to the learning plateau (trials 201-300). For all other subjects, the final errors were smaller than the initial errors (Figure 5A). There was a significant correlation between the initial and the final errors, meaning subjects with higher initial errors tended to have higher final errors as well.

**Figure 5.**
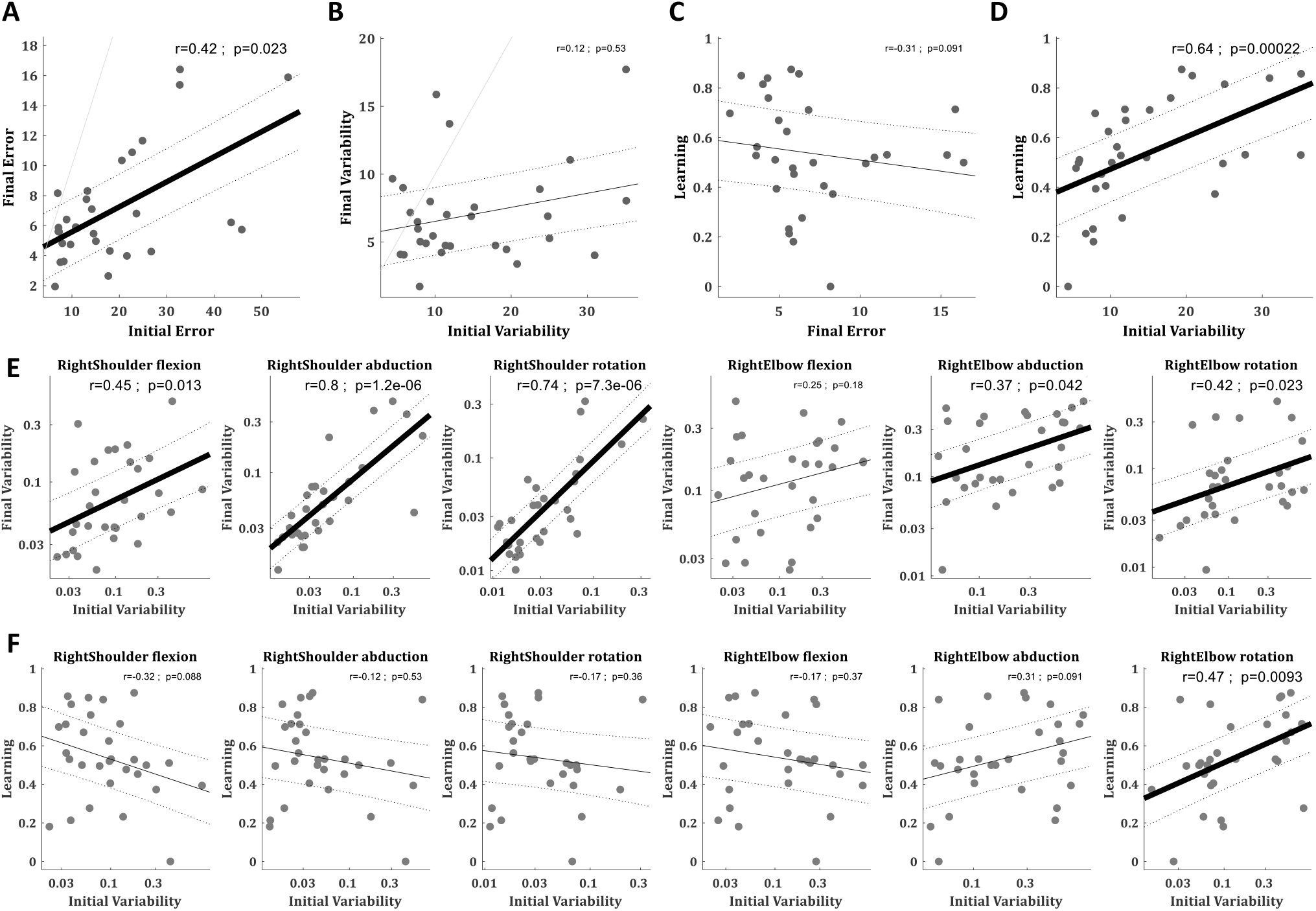
Variability and learning across subjects. (**A**) Correlation between subjects’ mean absolute directional error (in degrees) over the first block (trials 1-25) and the learning plateau (trials 201-300). (**B**) Correlation between subjects’ directional variability (in degrees) over first block (corrected for learning trend, see text) and over the learning plateau. (**C**) Correlation between subjects’ mean absolute directional error over the learning plateau and their learning. (**D**) Correlation between subjects’ directional variability over the first block (corrected for learning trend, see text) and their learning. (**E**) Correlation between subjects’ VPE variability (in logarithmic scale) over the first block and the learning plateau for the right arm joints. (**F**) Correlation between subjects’ VPE variability (in logarithmic scale) over the first block and their learning for the right arm joints. (**A-F**) Correlation values are Spearman rank correlation, p-values are FDR corrected for multiple comparisons, regression lines (black) are linear fits with 95% confidence intervals (doted lines). (**A, B**) unity lines are in grey.

While over the learning most subjects decreased their intertrial variability in the outcome (ball direction; Figure 1H & 5B) there was some tendency (though non-significant) for subjects who were initially more variable to be also more variable after learning (Figure 5B). The intertrial variability of the joint angular velocity profiles, which also decreased over learning (Figure 4B), showed a clearer and stronger correlation between the initial and the final intertrial variability (Figure 5E & Supplementary Figure 4). While this phenomenon was observed in various joints across the body, and dominant in the abduction across the spine joints, it was most dominant in the right shoulder abduction and rotation, the two joint angles that do most of the movement and carry most of its variance (Figure 2).

Learning was defined as the difference between the initial error (over the first block: trials 1-25) and the final error (over the learning plateau: trials 201-300) normalised by the initial error. For the one subject who showed no learning (had bigger errors during the learning plateau than during the first block), we set learning to zero to avoid negative learning value. While there is a negative relation between learning and final error by definition, due to the normalization by the initial error (which was highly variant), there was no significant correlation between the learning and the final error (as subjects who started worse could have learned more but still not perform better after learning), but there was only a trend that more learning leads to smaller final errors (Figure 5C). We speculated that the manipulative complexity (the degrees of freedom in the movement) might explain part of the inter-subject variability in learning rates. Presumably, subjects that learn more also show a higher increase in their manipulative complexity. Yet, we found no such relation. Both the initial (over the first block: trials 1-25) and the final (over the learning plateau: trials 201-300) manipulative complexity levels showed only a weak, non-significant, correlation to learning, and the increase in manipulative complexity showed no correlation to learning (Supplementary Figure 5).

We then tested if higher levels of initial task-relevant motor variability (variability in the directional error of the target ball) in this complex real-world task could predict faster learning across individuals, as found in simple lab experiments^38^. We indeed found that individuals with higher intertrial variability in the directional error of the target ball over the first block showed more learning (Spearman rank correlation r=0.64, p<0.001; Figure 5D). Importantly, this is the corrected intertrial variability (as in Figure 1H) which is calculated over the residuals from a regression line fitted to the ball direction to correct for the learning that is happening within the block. As a control, we also tested for correlation with the initial variability in the target ball velocity – which is a task-irrelevant motor variability – and found no correlation (Spearman rank correlation r=0.06, p=0.77).

Since much of the learning is happening during the first 25 trials, calculating learning over blocks can lead to a ceiling effect. Therefore, to test the robustness of the correlation between learning and variability, we re-calculated the learning rate based on the 5, 7, 10, 15, and 20 initial trials (Supplementary Figure 6). All choices of more than 5 trials showed a significant correlation between the initial variability and the learning. In the case of 5 trials, there were 2 outlier subjects with high variability and little learning who damaged the correlation. These 2 subjects had an initial ‘lucky shot’ which biased the learning calculation. Once including more trials, this effect was washed out.

Next, we tested the link between learning and initial variability over the joint angular velocity profiles of the right arm (Figure 5F). We found that the only joint angle where the intertrial variability showed a significant correlation to learning was the right elbow rotation (Spearman rank correlation r=0.47, p=0.0086), which is the forearm supination. We further tested the link over the full-body kinematics (Supplementary Figure 7) and found no other joint that showed this correlation. Thus, while learning leads to an overall reduction in movement variability, only initial variability in specific, task-relevant, dimensions can facilitate/predict learning.

## Discussion

In this paper, we introduce a new paradigm for studying naturalistic motor learning during whole-body movement in a complex real-world motor skill task. Our results present new insights into motor learning in the real-world. While the learning curves in this in-the-wild paradigm are within the same range of those reported in reductionistic motor adaptation tasks^2,39^ we find that this learning is taking place not only in the task-relevant joints but across the entire body. Also, we found that task-relevant initial variability in the ball direction (movement outcome) can predict learning, like in laboratory-tasks^38^, and so can the initial variability in the right forearm supination which is the task-relevant joint angle variability.

While pushing towards real-world neuroscience, we started here with a relatively constrained version of the real-world task, asking subjects to perform repeated trials of the same pool shot. This was to enable analysis using well-developed methods of laboratory-tasks. Nonetheless, it is a major step in the direction of a naturalistic study. First, we allow full-body unconstrained movement. Second, we do not use any artificial go cue and allow self-paced movement and as many preparatory movements as the subject needs for each shoot. Third, subjects receive natural somatosensory feedback. And last, we do not perturb the feedback to induce learning.

### Fundamentals of real-world motor learning

Across all subjects, we found that motor learning is a holistic process - the body is affected as a whole by learning the task. This was evident in the decrease in the VPE and the intertrial variability over learning (Figure 4A & B). This result should not come as a surprise considering decades of research in sport science showing this relationship. For example, baseball pitcher’s torso, pelvis, and leg movements are directly associated with ball velocity^40–42^. Recently it was also demonstrated with full-body motion capture in a ball throwing task^43^. And yet, unlike baseball pitches, basketball throws, or any unconstrained overarm throw, where the whole body is moving, in a pool shot the shooting arm is doing most of the movement and there is very little body movement. Thus, the whole-body learning is not trivial and suggestive that even in arm movement laboratory-tasks there is probably a whole-body learning aspect that is overlooked.

We also found a proximal-to-distal gradient in the learning rates over the right arm joints (Figure 4A & Supplementary Figure 3). This is especially interesting in light of the well-known phenomenon of proximal-to-distal sequence in limb movements in sports science^44^ and rehabilitation^45^. While there are records of proximal-to-distal sequence at multiple time scales^46^, our results are the first to suggest that this gradient also occur over repetitions as part of the learning process.

### Variability & learning

Intertrial variability is a fundamental characteristic of human movements and its underling neural activity^47^. It was recently reported that individuals exhibit distinct magnitudes of movement variability, which are consistent across movements and effectors, suggesting individual traits in movement variability^48^. Our results show that subjects who were initially more variable tended to be also more variable after learning in many joints across the body (Figure 5E & Supplementary Figure 4) and specifically in those of right shoulder that carry most of the variance in the movement. This result is in-line with the notion that there is an individual trait in movement variability.

Intertrial kinematic variability is also thought to be critical for motor learning^49–53^. It was suggested that individuals with higher levels of task-relevant movement variability exhibit faster motor learning in both skill learning and motor adaptation error-based paradigms^38^. The failures to reproduce this result in visuomotor adaptation studies^54,55^, led to the idea that experiments with task-relevant feedback (which is common in visuomotor studies) emphasize execution noise over planning noise, whereas measurements made without feedback (as in^38^) may primarily reflect planning noise^53^. This is in-line with a recent modelling work in a visuomotor adaptation study (with task-relevant feedback) in which subjects with higher planning noise showed faster learning, but the overall movement variability was dominated by execution noise that was negatively correlated with learning^56^. In our task there were no manipulations or perturbations, thus, task-relevant feedback was fully available to the participants. On the other hand, in real-world, there is no baseline variability, and the variability was measured during early learning and therefore is probably dominated by planning noise, as subjects explore, regardless of the visual feedback. Indeed, subjects with higher variability in the target ball direction over the first block showed higher learning rates (Figure 5D). Our results straighten the link between variability and learning and are the first to show that it applies to real-world tasks. Moreover, the only joint angle that showed a significant correlation between initial variability and learning was the right forearm supination (measured by the right elbow rotation in our IMUs setup, Figure 5F & Supplementary Figure 7). Following the idea that task-relevant variability predicts learning, it would suggest that the right elbow rotation is the task-relevant joint angle to adjust during the initial learning of a simple pool shoot. Indeed, guidebooks for pool and billiards emphasize that while shooting one should keep one’s body still and move only the back (right) arm from the elbow down. While the elbow flexion movement gives the power to the shoot, the forearm supination (also known as ‘screwing’ in billiards) maintains the direction of the cue.

It is important to note that this refers specifically to the forearm supination around the elbow and not around the wrist. This is due to the nature of the data collected with the sensors suit where the joint angles are recorded with 3 degrees of freedom based on the angles between the sensors from both sides of each joint. Thus, hinge joints of the body which have only one anatomical degree of freedom been recorded as 3 Euler angles. Specifically, the elbow rotation is the rotation between the upper arm sensor and the forearm sensor and is equivalent to forearm supination around the elbow. The wrist rotation is the rotation between the forearm sensor and the hand sensor and is equivalent to hand supination.

It is also important to highlight that the above are correlational and cannot address the question of causality: i.e. can higher initial variability cause faster learning? While the study of real-world tasks takes us closer to understanding real-world motor-learning, it is lacking the key advantage of laboratory tasks (which made them so popular) of highly controlled manipulations of known variables, to isolate specific movement/learning components. To address this issue and introduce manipulations to this real-world task and establish causality, we developed an embodied virtual reality version of our pool task^57^. The VR-based approach can overcome this limitation.

## Conclusions

In this study, we demonstrate the feasibility and importance of studying human neuroscience in-the-wild, and specifically in naturalistic real-world skill tasks. While finding similarities in learning structure between our real-world paradigm and lab-based motor learning studies, we highlight crucial differences, namely, real-world motor learning is a holistic full-body process. Looking at the motor behaviour over learning across the entire body enabled us to explore the relationship between variability and learning and define task-relevant variability that can facilitate learning.

## Methods

### Ethics statement

All experimental procedures were approved by Imperial College Research Ethics Committee and performed in accordance with the declaration of Helsinki. All subjects gave informed consent prior to participating in the study.

### Experimental Setup and Design

30 right-handed healthy human volunteers with normal or corrected-to-normal visual acuity (12 women and 18 men, aged 24±3) participated in the study. The recruitment criteria were that they played pool/billiards/snooker for leisure fewer than 5 times in their life, never in the recent 6 months, and had never received any pool game instructions. All volunteers gave informed consent before participating in the study, and all experimental procedures were approved by the Imperial College Research Ethics Committee and performed in accordance with the declaration of Helsinki. The volunteers stood in front of a 5ft pool table (Riley Leisure, Bristol, UK) with 1 7/8” (48mm diameter) pool balls. Volunteers performed 300 repeated trials where the cue ball (white) and the target ball (red) were placed in the same locations. We asked volunteers to shoot the target ball towards the pocket of the far-left corner (Figure 1A). Trials were split into 6 sets of 50 trials with a short break in-between to allow the subjects to rest a bit and reduce potential fatigue. Each experimental set (of 50 trials) took 8 to 12 minutes. For the data analysis, we further split each set into two blocks of 25 trials each, resulting in 12 blocks. During the entire learning process, we recorded the subjects’ full-body movements with a motion-tracking ‘suit’ of 17 wireless inertial measurement units (IMUs; Figure 1B). The balls on the pool table were tracked with a high-speed camera (Dalsa Genie Nano, Teledyne DALSA, Waterloo, Ontario) to assess the subjects’ success in the game and to analyze the changes throughout learning, not only in the body movement but also in its outcome - the ball movement (Figure 1C).

### Balls tracking

The balls movement on the pool table were tracked with a computer vision system mounted from the ceiling. The computer vision camera was a Genie Nano C1280 Color Camera (Teledyne Dalsa, Waterloo, Canada), colour images were recorded with a resolution of 752×444 pixels and a frequency of 200Hz. This Ethernet-based camera was controlled via the Common Vision Blox Management Console (Stemmer Imaging, Puchheim, Germany) and image videos recorded with our custom software written in C++ based on a template provided by Stemmer Imaging. Our software captured the high-performance event timer, the camera frames and converted the images from the camera’s proprietary CVB format to the open-source OpenCV (https://opencv.org/) image format for further processing in OpenCV. The video frames were stored as an uncompressed AVI file to preserve the mapping between pixel changes and timings and the computer’s real-time clock time-stamps were recorded to a text file. Each trial was subject-paced, so the experimenter observed the subject and hit the spacebar key as an additional trigger event to the time-stamps text file. This timing data was later used to assist segmentation of the continuous data stream into trials. The positions of the two pool balls (white cue ball and red target ball) were calculated from the video recordings offline using custom software written in C++ using OpenCV. Then, with custom software written in MATLAB (R2017a, The MathWorks, Inc., MA, USA), we segmented the ball tracking data and extracted the trajectory of the balls in each trial. For each trial, a 20 × 20 pixels (approx 40 x 40 mm) bounding box was set around the centre of the 48 mm diameter cue ball. The time the centre of the ball left the bounding box was recorded as the beginning of the cue ball movement. The pixel resolution and frame rate were thus sufficient to detect movement onset, acceleration and deceleration of the pool balls. The target (red) ball initial position and its position in the point of its peak velocity were used to calculate the ball movement angle (relative to a perfectly straight line between the white cue ball and the red target ball). We subtracted this angle from the centre of the pocket angle (the angle the target ball initial position and the centre of the pocket relative to the same straight line between the balls) to calculate the directional error for each shot.

### Full-Body Motion Tracking

Kinematic data were recorded at 60 Hz using a wearable motion tracking ‘suit’ of 17 wireless IMUs (Xsens MVN Awinda, Xsens Technologies BV, Enschede, The Netherlands). Data acquisition was controlled via a graphical interface (MVN Analyze, Xsens Technologies BV, Enschede, The Netherlands). Xsens MVN uses a biomechanical model and proprietary algorithms to estimate 3D joint kinematics ^58,59^. The Xsens sensors shows high accuracy^34^, and the Xsens MVN system was used and validated in tracking real-world behaviour in many sports including football^60^, horse riding^61^, ski^62^ and snowboarding^63^. The Xsens 3D joint kinematics were exported as XML files and analysed using custom software written in MATLAB (R2017a, The MathWorks, Inc., MA, USA). The Xsens full-body kinematics were extracted in joint angles in 3 degrees of freedom for each joint that followed the International Society of Biomechanics (ISB) recommendations for Euler angle extractions of Z (flexion/extension), X (abduction/adduction) Y (internal/external rotation). This standard approach includes hinge joints of the body which have only 1 degree of freedom being recorded as 3 Euler angles.

### Angular Velocity Profile Analysis

From the Xsens 3D joint angles we extracted the angular velocity profiles of all joints in all trials. We defined the peak of the trial as the peak of the average absolute angular velocity across the DoFs of the right shoulder and the right elbow. We aligned all trials around the peak angular velocity of the trial and cropped a window of 1 sec around the peak for the analysis of joint angular velocity profiles during the shot and its follow-through. This time window covered the entire movement of the pool shoot while eliminating the preparatory movement and the mock shoots (Figure 2).

### Task performance & learning measures

The task performance was measured by the trial error which was defined as an absolute angular difference between the target ball movement vector direction and the desired direction to land the target ball in the centre of the pocket. The decay of error over trials is the clearest signature of learning in the task. For measuring success rates and intertrial variability we divided the trials into blocks of 25 trials by dividing each experimental set of 50 trials to two blocks. This was done to increase the resolution in time from calculating those on the full sets. Success rate in each block was defined by the ratio of successful trial (in which the ball fell into the pocket). To improve robustness and account for outliers, we fitted the errors in each block with a t-distribution and used the location and scale parameters (μ and σ) as the blocks’ centre and variability measures. To correct for learning within a block, we also calculated a corrected intertrial variability, which was the intertrial variability over the residuals from a regression line fitted to the ball direction in each block. This correction for the learning trend within a block does not change the variability measure by much (Figure 1G&H). This is since our variability measure is not the standard deviation, but the scale parameter of a t-distribution fitted to the errors. When correcting the change in the distribution fitted was mostly an increase in the degrees of freedom and a not decrease in the scale. I.e. the early trials which were much higher than the mean and the late trials which were much lower become closer to the mean and therefore the distribution is more normal as it loses the heavy tails). This is highlighting the robustness of the scale measure for variability.

To quantify the within-trial variability structure of the body movement, we use the generalised variance, which is the determinant of the covariance matrix^35^ and is intuitively related to the multidimensional scatter of data points around their mean. We measured the generalised variance over the velocity profiles of all joints in each trial to see how it changes with learning. To study the complexity of the body movement which was defined by the number of degrees of freedom used by the subject we applied principal component analysis (PCA) across joints for the velocity profiles per trial for each subject and used the number of PCs that explain more than 1% of the variance to quantify the degrees of freedom in each trial movement. We also calculated the manipulative complexity which was suggested by Belić and Faisal^37^ as a way to quantify complexity for a given number of PCs on a fixed scale (C = 1 implies that all PCs contribute equally, and C = 0 if one PC explains all data variability).

### Statistical Analysis

Trial by trial learning curves of single-trial performance measure (directional error of the target ball relative to the centre of the pocket) were fitted with a single, double, and triple exponential learning curve using Matlab fit function. As in most motor learning datasets, the double exponential curve showed the best fit (Supplementary Figure 1).

As a measure of task performance in body space, correlation distances (one minus Pearson correlation coefficient) were calculated between the angular velocity profile of each joint in each trial to the angular velocity profiles of that joint in all successful trials. The minimum over these correlation distances produced a single measure of Velocity Profile Error (VPE) for each joint in each trial.

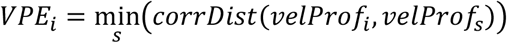

Thus, VPE in trial *i* was the minimal correlation distances between the angular velocity profile in trial *i* (*velProf*_*i*_) and the angular velocity profiles in successful trials *s* (*velProf*_*s*_). While there are multiple combinations of body variables that can all lead to successful task performance, this measure looks for the distance from the nearest successful solution used by the subjects and thus provides a metric that accounts for the redundancy in the body.

All correlations between error, variability, and learning are Spearman’s rank correlation coefficients to be robust to outliers and non-linear trends, and their p-values are FDR corrected for multiple comparisons. Regression lines are based on linear regression fits (in logarithmic scale for VPE variability) and are presented with 95% confidence intervals.

## Acknowledgements

We thank Marlene Gonzalez for her contribution to the data collection and our participants for taking part in the study. We acknowledge the technical and computational support of Alex Harston and Chaiyawan Auepanwiriyakul. We thank Alex Harston for helpful comments on the manuscript. The study was enabled by financial support to a Royal Society-Kohn International Fellowship (NF170650; SH & AAF) and by eNHANCE (http://www.enhance-motion.eu) under the European Union’s Horizon 2020 research and innovation program grant agreement No. 644000 (SH & AAF).

## Notes

### Competing Interest Statement

The authors have declared no competing interest.

## References

1. Shadmehr, R. & Mussa-Ivaldi, F. A. Adaptive representation of dynamics during learning of a motor task. J. Neurosci. 14, 3208–24 (1994).

2. Smith, M. A., Ghazizadeh, A. & Shadmehr, R. Interacting adaptive processes with different timescales underlie short-term motor learning. PLoS Biol. 4, e179 (2006).

3. Diedrichsen, J., Hashambhoy, Y., Rane, T. & Shadmehr, R. Neural correlates of reach errors. J. Neurosci. 25, 9919–9931 (2005).

4. Howard, I. S., Wolpert, D. M. & Franklin, D. W. The Value of the Follow-Through Derives from Motor Learning Depending on Future Actions. Curr. Biol. 25, 397–401 (2015).

5. Krakauer, J. W., Pine, Z., Ghilardi, M. & Ghez, C. Learning of visuomotor transformations for vectorial planning of reaching trajectories. J. Neurosci. 20, 8916–8924 (2000).

6. Mazzoni, P. & Krakauer, J. An Implicit Plan Overrides an Explicit Strategy during Visuomotor Adaptation. J. Neurosci. 26, 3642–3645 (2006).

7. Taylor, J. a, Krakauer, J. W. & Ivry, R. B. Explicit and implicit contributions to learning in a sensorimotor adaptation task. J. Neurosci. 34, 3023–32 (2014).

8. Haar, S., Donchin, O. & Dinstein, I. Dissociating Visual and Motor Directional Selectivity Using Visuomotor Adaptation. J. Neurosci. 35, 6813–6821 (2015).

9. Bromberg, Z., Donchin, O. & Haar, S. Eye movements during visuomotor adaptation represent only part of the explicit learning. eNeuro 6, 1–12 (2019).

10. Reis, J. et al. Noninvasive cortical stimulation enhances motor skill acquisition over multiple days through an effect on consolidation. Proc. Natl. Acad. Sci. U. S. A. 106, 1590–5 (2009).

11. Ma, L., Narayana, S., Robin, D. A., Fox, P. T. & Xiong, J. Changes occur in resting state network of motor system during 4weeks of motor skill learning. Neuroimage 58, 226–233 (2011).

12. Clerget, E., Poncin, W., Fadiga, L. & Olivier, E. Role of Broca’s Area in Implicit Motor Skill Learning: Evidence from Continuous Theta-burst Magnetic Stimulation. J. Cogn. Neurosci. 24, 80–92 (2012).

13. Yokoi, A., Arbuckle, S. A. & Diedrichsen, J. The role of human primary motor cortex in the production of skilled finger sequences. J. Neurosci. 38, 1430–42 (2018).

14. Ingram, J. N. & Wolpert, D. M. Naturalistic approaches to sensorimotor control. Prog. Brain Res. 191, 3–29 (2011).

15. Wolpert, D. M., Diedrichsen, J. & Flanagan, J. R. Principles of sensorimotor learning. Nat. Rev. Neurosci. 12, 739–51 (2011).

16. Faisal, A., Stout, D., Apel, J. & Bradley, B. The Manipulative Complexity of Lower Paleolithic Stone Toolmaking. PLoS One 5, e13718 (2010).

17. Hecht, E. E. et al. Acquisition of Paleolithic toolmaking abilities involves structural remodeling to inferior frontoparietal regions. Brain Struct. Funct. 220, 2315–2331 (2014).

18. Laughlin, S. A simple coding procedure enhances a neuron’s information capacity. Zeitschrift fur Naturforschung - Section C Journal of Biosciences 36, 910–912 (1981).

19. Simoncelli, E. P. & Olshausen, B. A. Natural Image Statistics and Neural Representation. Annu. Rev. Neurosci. 24, 1193–1216 (2001).

20. Cohen, R. G. & Sternad, D. Variability in motor learning: relocating, channeling and reducing noise. Exp. Brain Res. 193, 69–83 (2009).

21. Abe, M. O. & Sternad, D. Directionality in distribution and temporal structure of variability in skill acquisition. Front. Hum. Neurosci. 7, 225 (2013).

22. Shmuelof, L., Krakauer, J. W. & Mazzoni, P. How is a motor skill learned? Change and invariance at the levels of task success and trajectory control. J. Neurophysiol. 108, 578–594 (2012).

23. Yang, J. F. & Scholz, J. P. Learning a throwing task is associated with differential changes in the use of motor abundance. Exp. Brain Res. 163, 137–158 (2005).

24. Scholz, J., Klein, M. C., Behrens, T. E. & Johansen-Berg, H. Training induces changes in white-matter architecture. Nat Neurosci 12, 1370–1371 (2009).

25. Sampaio-Baptista, C. et al. Gray matter volume is associated with rate of subsequent skill learning after a long term training intervention. Neuroimage 96, 158–166 (2014).

26. Sampaio-Baptista, C. et al. Changes in functional connectivity and GABA levels with long-term motor learning. Neuroimage 106, 15–20 (2015).

27. Ono, Y. et al. Motor learning and modulation of prefrontal cortex: an fNIRS assessment. J. Neural Eng. 12, 066004 (2015).

28. Haar, S. & Faisal, A. A. Brain activity reveals multiple motor-learning mechanisms in a real-world task. Front. Hum. Neurosci. (2020). doi:10.3389/FNHUM.2020.00354

29. Thomik, A. A. C. On the structure of natural human movement. (Imperial College London, 2016).

30. Phelan, M. The Game of Billiards. (D. Appleton and Company, New York, 1859).

31. De Vere, A. Billiards made easy, by ‘Winning Hazard’. (Houlston and Sons, London, 1873).

32. Mizerak, S. Pocket Billiards Tips and Trick Shots. (McGraw-Hill, 1982).

33. Leider, N. Pool & billiards for dummies. (Wiley Publishing, 2010).

34. Auepanwiriyakul, C., Waibel, S., Songa, J., Bentley, P. & Faisal, A. A. Accuracy and Acceptability of Wearable Motion Tracking Smartwatches for Inpatient Monitoring. medRxiv 2020.07.24.20160663 (2020). doi:10.1101/2020.07.24.20160663

35. Wilks, S. S. Certain Generalizations in the Analysis of Variance. Biometrika 24, 471 (1932).

36. Bernstein, N. The co-ordination and regulation of movements. (Pergamon Press, 1967).

37. Belić, J. J. & Faisal, A. A. Decoding of human hand actions to handle missing limbs in neuroprosthetics. Front. Comput. Neurosci. 9, 27 (2015).

38. Wu, H. G., Miyamoto, Y. R., Gonzales Castro, L. N., Ölveczky, B. C. & Smith, M. A. Temporal structure of motor vriability is dynamically regulated and predicts motor learning ability. Nat. Neurosci. 17, 312–321 (2014).

39. McDougle, S. D., Bond, K. M. & Taylor, J. A. Explicit and Implicit Processes Constitute the Fast and Slow Processes of Sensorimotor Learning. J. Neurosci. 35, 9568–9579 (2015).

40. Kageyama, M., Sugiyama, T., Takai, Y., Kanehisa, H. & Maeda, A. Kinematic and Kinetic Profiles of Trunk and Lower Limbs during Baseball Pitching in Collegiate Pitchers. J. Sports Sci. Med. 13, 742–50 (2014).

41. Oliver, G. D. & Keeley, D. W. Pelvis and torso kinematics and their relationship to shoulder kinematics in high-school baseball pitchers. J. Strength Cond. Res. 24, 3241–3246 (2010).

42. Stodden, D. F., Langendorfer, S. J., Fleisig, G. S. & Andrews, J. R. Kinematic Constraints Associated With the Acquisition of Overarm Throwing Part I. Res. Q. Exerc. Sport 77, 417–427 (2006).

43. Maselli, A. et al. Where Are You Throwing the Ball? I Better Watch Your Body, Not Just Your Arm! Front. Hum. Neurosci. 11, 505 (2017).

44. Herring, R. M. & Chapman, A. E. Effects of changes in segmental values and timing of both torque and torque reversal in simulated throws. J. Biomech. 25, 1173–1184 (1992).

45. Twitchell, T. E. The restoration of motor function following hemiplegia in man. Brain 74, 443–480 (1951).

46. Serrien, B. & Baeyens, J. P. The proximal-to-distal sequence in upper-limb motions on multiple levels and time scales. Hum. Mov. Sci. (2017). doi:10.1016/j.humov.2017.08.009

47. Faisal, a A., Selen, L. P. J. & Wolpert, D. M. Noise in the nervous system. Nat. Rev. Neurosci. 9, 292–303 (2008).

48. Haar, S., Donchin, O. & Dinstein, I. Individual Movement Variability Magnitudes Are Explained by Cortical Neural Variability. J. Neurosci. 37, 9076–9085 (2017).

49. Herzfeld, D. J. & Shadmehr, R. Motor variability is not noise, but grist for the learning mill. Nat. Neurosci. 17, 149–50 (2014).

50. Teo, J. T. H., Swayne, O. B. C., Cheeran, B., Greenwood, R. J. & Rothwell, J. C. Human theta burst stimulation enhances subsequent motor learning and increases performance variability. Cereb. Cortex 21, 1627–1638 (2011).

51. Braun, D. a, Aertsen, A., Wolpert, D. M. & Mehring, C. Motor Task Variation Induces Structural Learning. Curr. Biol. 19, 352–357 (2009).

52. Wilson, C., Simpson, S. E., van Emmerik, R. E. a & Hamill, J. Coordination variability and skill development in expert triple jumpers. Sports Biomech. 7, 2–9 (2008).

53. Dhawale, A. K., Smith, M. A. & Ölveczky, B. P. The Role of Variability in Motor Learning. Annu. Rev. Neurosci. 40, 479–498 (2017).

54. Singh, P., Jana, S., Ghosal, A. & Murthy, A. Exploration of joint redundancy but not task space variability facilitates supervised motor learning. Proc. Natl. Acad. Sci. 113, 14414–14419 (2016).

55. He, K. et al. The Statistical Determinants of the Speed of Motor Learning. PLOS Comput. Biol. 12, e1005023 (2016).

56. van der Vliet, R. et al. Individual Differences in Motor Noise and Adaptation Rate Are Optimally Related. eneuro 5, ENEURO.0170–18.2018 (2018).

57. Haar, S., Sundar, G. & Faisal, A. A. Embodied virtual reality for the study of real-world motor learning. bioRxiv 1–14 (2020). doi:10.1101/2020.03.19.998476

58. Roetenberg, D., Luinge, H. & Slycke, P. Xsens MVN: full 6DOF human motion tracking using miniature inertial sensors. Xsens Motion Technol. BV, … 8, 1–7 (2009).

59. Schepers, M., Giuberti, M. & Bellusci, G. Xsens MVN : Consistent Tracking of Human Motion Using Inertial Sensing. Xsens Technol. 1–8 (2018). doi:10.13140/RG.2.2.22099.07205

60. Blair, S., Duthie, G., Robertson, S., Hopkins, W. & Ball, K. Concurrent validation of an inertial measurement system to quantify kicking biomechanics in four football codes. J. Biomech. 73, 24–32 (2018).

61. Gandy, E. A., Bondi, A., Hogg, R. & Pigott, T. M. C. A preliminary investigation of the use of inertial sensing technology for the measurement of hip rotation asymmetry in horse riders. Sport. Technol. 7, 79–88 (2014).

62. Lee, S. K., Kim, K., Kim, Y. H. & Lee, S. S. Motion anlaysis in lower extremity joints during ski carving turns using wearble inertial sensors and plantar pressure sensors. in 2017 IEEE International Conference on Systems, Man, and Cybernetics, SMC 2017 2017-January, 695–698 (Institute of Electrical and Electronics Engineers Inc., 2017).

63. Krüger, A. & Edelmann-Nusser, J. Biomechanical analysis in freestyle snowboarding: Application of a full-body inertial measurement system and a bilateral insole measurement system. Leis. Loisir 2, 17–23 (2009).

