## Supplementary figures for "Motor learning in real-world pool billiards"

### Supplementary Information

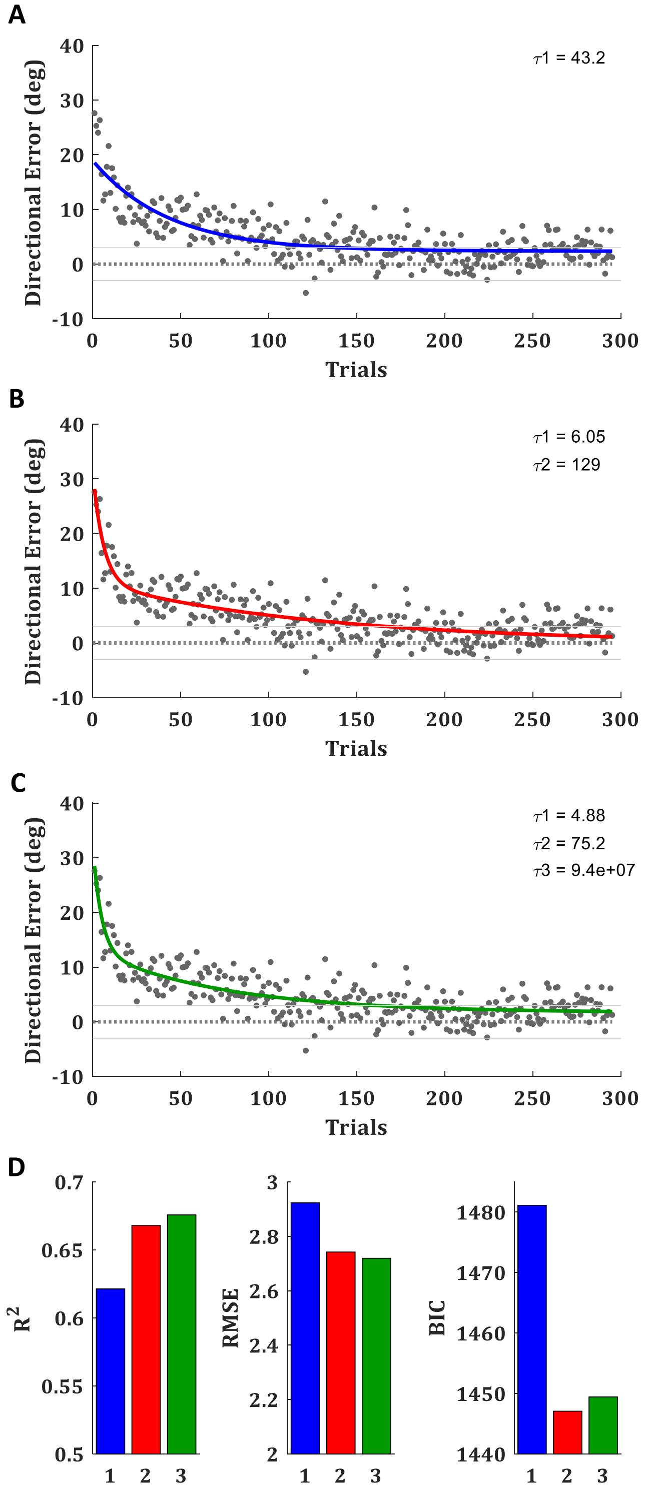


**Supplementary Figure 1.** *Exponential fits*. (**A-C**) The trial-by-trial directional error of the target-ball (relative to the direction from its origin to the centre of the target pocket), averaged across all subjects, with (**A**) single, (**B**) double, and (**C**) triple exponential fit (blue, red, and green curves, respectively). Grey lines mark the range of successful trials (less than 3 degrees form the centre of the pocket). (**D**) The R square ($R^2$), root mean square error (SME), and Bayesian information criterion (BIC) for the 3 fits.

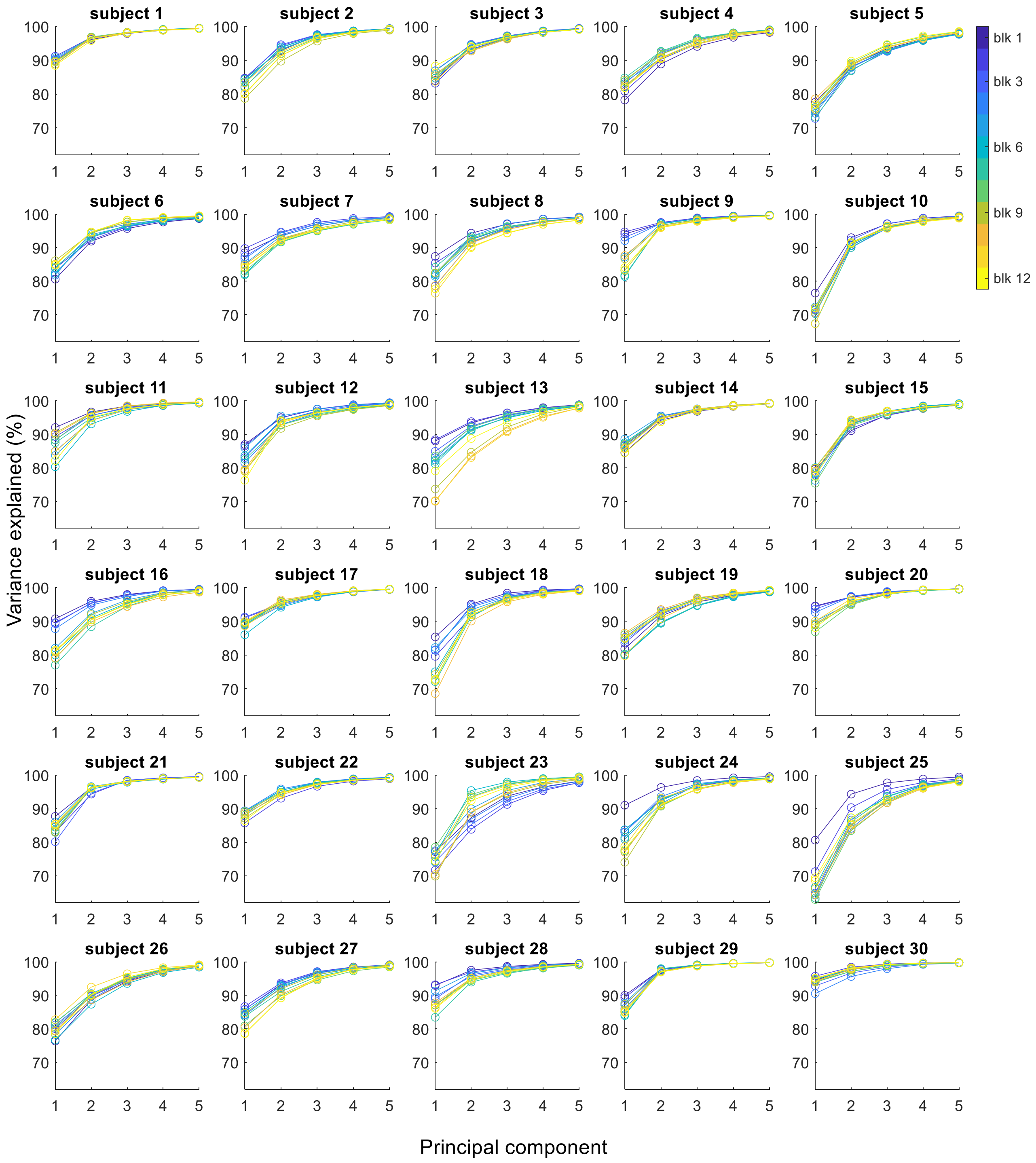


**Supplementary Figure 2.** *PCA variance explained*. The variance explained by the first five principal components for each subject in each block of trials. Lines are averaged over all trials within each block. Colour-code is by blocks order, from blue in the first block (blk 1) to yellow in the last block (blk 12).

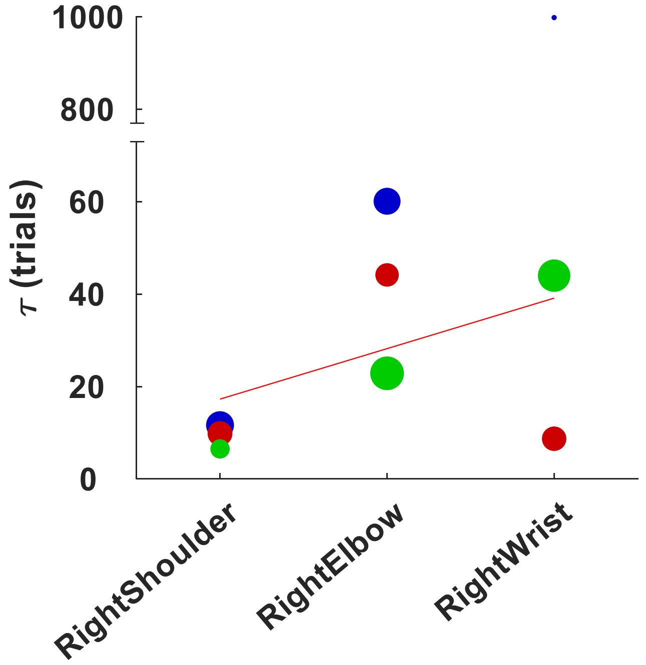


**Supplementary Figure 3.** *Proximal-to-distal gradient in the VPE over the right arm joints*. The time constants of the exponential fits of the trial-by-trial Velocity Profile Error for all 3 DoF of the right arm joints (from figure 4A). The size of the dot represents the learning rate (B in Supplementary Table 1) and the colour code is the same as in figure 4 (blue: flexion/extension; red: abduction/adduction; green: internal/external rotation). The ret line is weighted linear fit, where the wight is the learning rate.

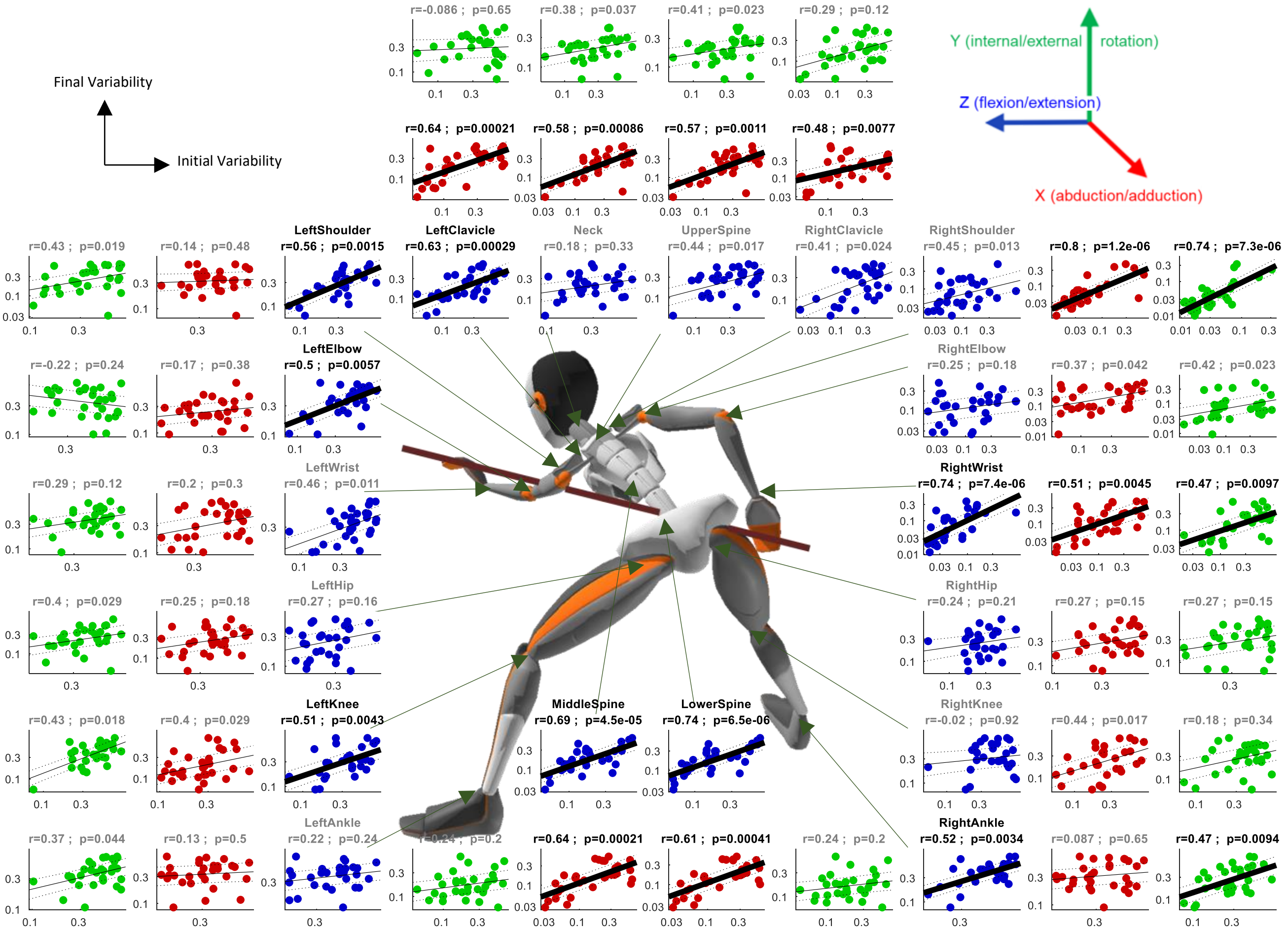


**Supplementary Figure 4.** *Correlation between subjects' VPE variability over the first block and over the learning plateau*. Presented for all joints in 3 degrees of freedom (DoF) for each joint (blue: flexion/extension, red: abduction/adduction; green: internal/external rotation). Subjects' VPE variability is in logarithmic scale. Correlation values are Spearman rank correlation, regression lines (black) are linear fits with 95% confidence intervals (doted lines).

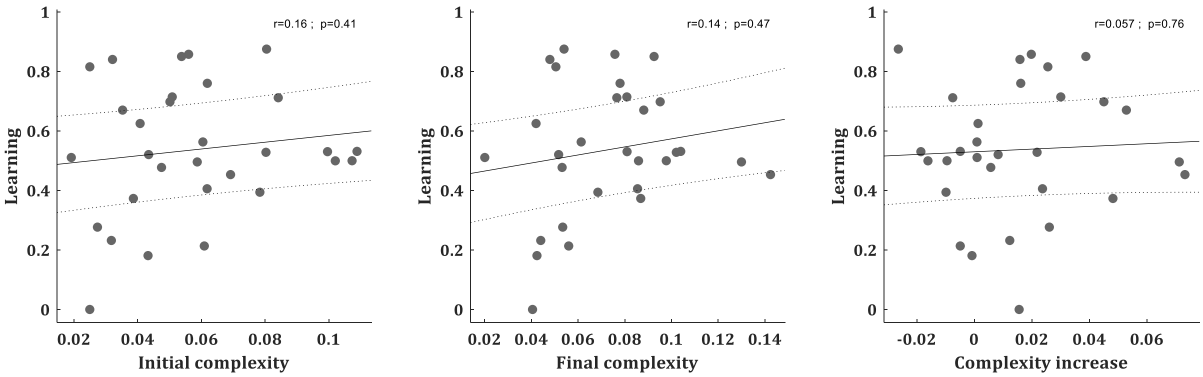


**Supplementary Figure 5.** *Manipulative complexity and learning across subjects*. (**A**) Correlation between subjects' manipulative complexity over the first block (trials 1-25) and their learning rates. (**B**) Correlation between subjects' manipulative complexity over the learning plateau (trials 201-300) and their learning rates. (**C**) Correlation between subjects' increase in manipulative complexity (from the first block to the learning plateau) and their learning rates. (**A-C**) Correlation values are Spearman rank correlation, regression lines (black) are linear fits with 95% confidence intervals (doted lines).

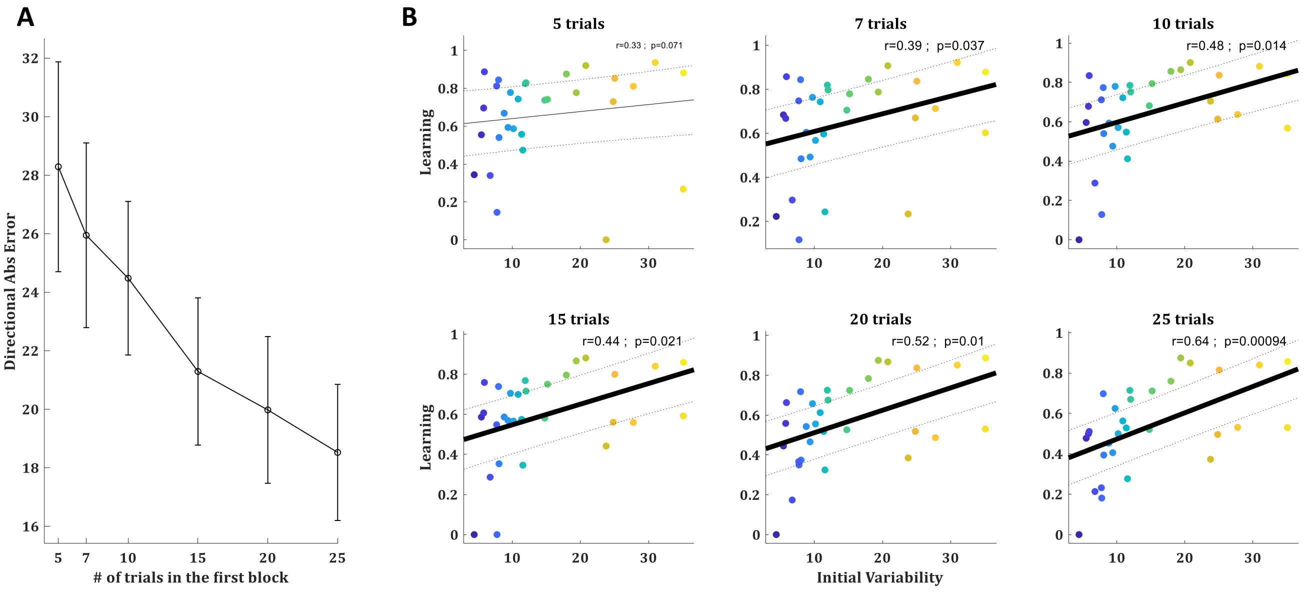


**Supplementary Figure 6.** *Variability and learning across subjects*. (**A**) The mean absolute directional error of the target-ball over the initial 5,7,10,15,20, and 25 trials, averaged across all subjects. Error bars represent SEM. (**B**) Correlation between subjects' directional variability over the first block (corrected for learning trend, see text) and their learning calculated based on their errors in the first 5,7,10,15,20, or 25 trials respectively in the different panels. Correlation values are Spearman rank correlation, p-values are FDR corrected for multiple comparisons, regression lines (black) are linear fits with 95% confidence intervals (doted lines).

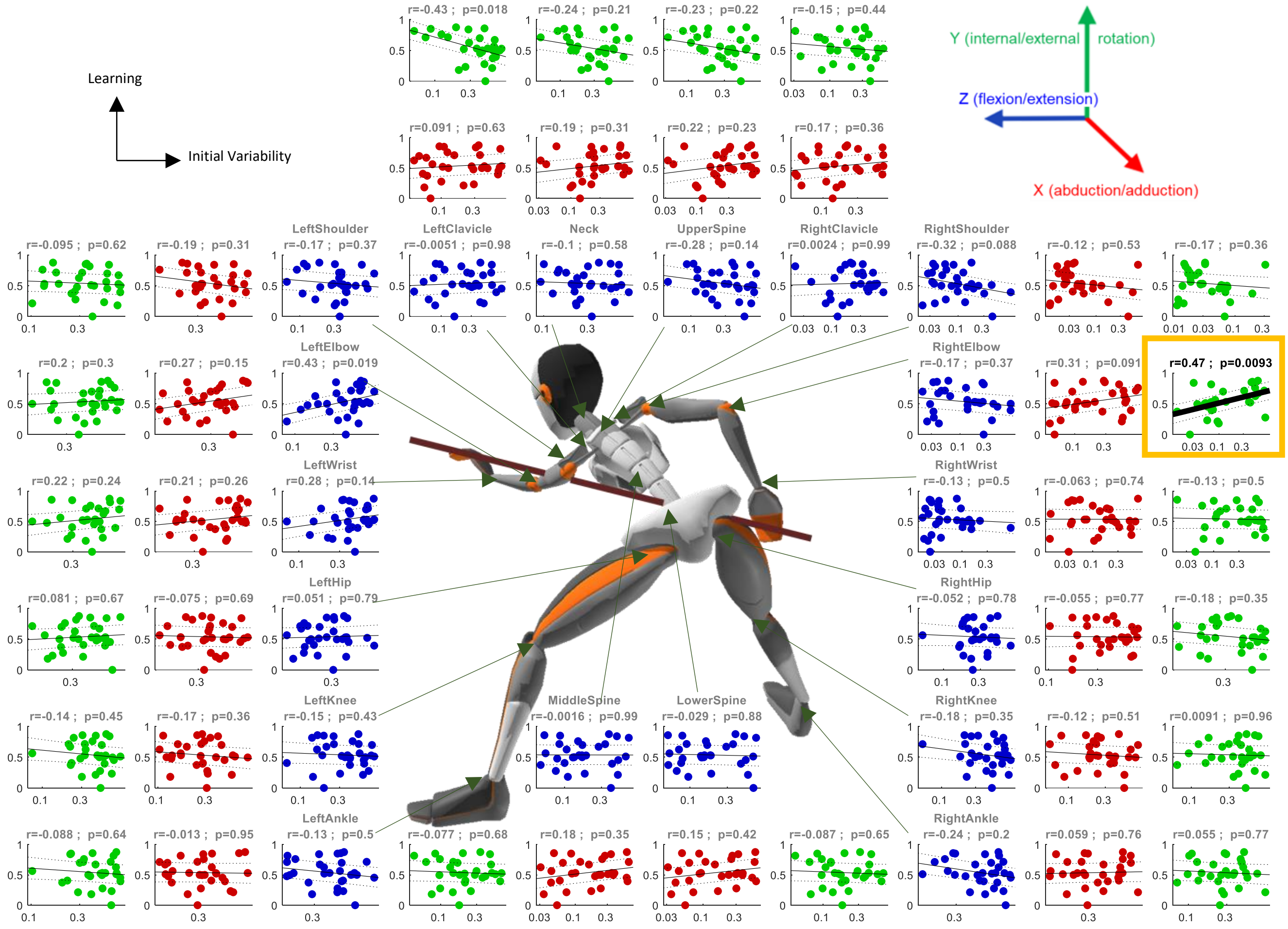


**Supplementary Figure 7.** *Correlation between subjects' VPE variability over first block and their learning*. Presented for all joints in 3 degrees of freedom (DoF) for each joint (blue: flexion/extension, red: abduction/adduction; green: internal/external rotation). Subjects' VPE variability is in logarithmic scale. Correlation values are Spearman rank correlation, regression lines (black) are linear fits with 95% confidence intervals (doted lines).

**Supplementary Table 1**. *Exponential fit parameters and Goodness of fit for the VPE (figure 4A)*

| jointsLabels | tau | B | R^2 | RMSE |
|---|---|---|---|---|
| LowerSpine flexion | 12.16 | 0.062 | 0.37 | 0.011 |
| LowerSpine abduction | 211.47 | 0.037 | 0.28 | 0.013 |
| LowerSpine rotation | 12.08 | 0.067 | 0.38 | 0.011 |
| MiddleSpine flexion | 11.58 | 0.057 | 0.33 | 0.011 |
| MiddleSpine abduction | 214.08 | 0.038 | 0.28 | 0.013 |
| MiddleSpine rotation | 10.86 | 0.075 | 0.39 | 0.012 |
| UpperSpine flexion | 16.92 | 0.100 | 0.55 | 0.014 |
| UpperSpine abduction | 15.74 | 0.099 | 0.56 | 0.013 |
| UpperSpine rotation | 33.80 | 0.094 | 0.64 | 0.015 |
| Neck flexion | 45.07 | 0.065 | 0.54 | 0.014 |
| Neck abduction | 15.45 | 0.099 | 0.52 | 0.014 |
| Neck rotation | 26.77 | 0.108 | 0.68 | 0.014 |
| RightClavicle flexion | 55.13 | 0.045 | 0.44 | 0.012 |
| RightClavicle abduction | 27.34 | 0.073 | 0.57 | 0.012 |
| RightClavicle rotation | 12.77 | 0.053 | 0.30 | 0.011 |
| RightShoulder flexion | 11.64 | 0.067 | 0.52 | 0.008 |
| RightShoulder abduction | 9.81 | 0.053 | 0.39 | 0.008 |
| RightShoulder rotation | 6.51 | 0.031 | 0.19 | 0.006 |
| RightElbow flexion | 60.08 | 0.063 | 0.59 | 0.013 |
| RightElbow abduction | 44.14 | 0.047 | 0.46 | 0.012 |
| RightElbow rotation | 22.86 | 0.099 | 0.72 | 0.011 |
| RightWrist flexion | 998.23 | 0.000 | 0.00 | 0.009 |
| RightWrist abduction | 8.71 | 0.051 | 0.24 | 0.010 |
| RightWrist rotation | 43.99 | 0.092 | 0.71 | 0.013 |
| LeftClavicle flexion | 34.92 | 0.061 | 0.46 | 0.014 |
| LeftClavicle abduction | 52.78 | 0.034 | 0.26 | 0.013 |
| LeftClavicle rotation | 50.67 | 0.057 | 0.44 | 0.015 |
| LeftShoulder flexion | 31.95 | 0.061 | 0.48 | 0.013 |
| LeftShoulder abduction | 21.34 | 0.073 | 0.45 | 0.014 |
| LeftShoulder rotation | 19.99 | 0.087 | 0.50 | 0.015 |
| LeftElbow flexion | 23.16 | 0.086 | 0.46 | 0.017 |
| LeftElbow abduction | 43.51 | 0.072 | 0.58 | 0.014 |
| LeftElbow rotation | 35.83 | 0.088 | 0.50 | 0.019 |
| LeftWrist flexion | 40.01 | 0.086 | 0.56 | 0.017 |
| LeftWrist abduction | 30.07 | 0.111 | 0.63 | 0.017 |
| LeftWrist rotation | 12.93 | 0.098 | 0.36 | 0.018 |
| RightHip flexion | 14.19 | 0.062 | 0.39 | 0.011 |
| RightHip abduction | 54.49 | 0.063 | 0.50 | 0.015 |
| RightHip rotation | 19.83 | 0.097 | 0.55 | 0.015 |
| RightKnee flexion | 48.79 | 0.112 | 0.67 | 0.018 |
| RightKnee abduction | 25.79 | 0.046 | 0.33 | 0.012 |
| RightKnee rotation | 39.95 | 0.096 | 0.65 | 0.015 |
| RightAnkle flexion | 40.10 | 0.114 | 0.72 | 0.016 |
| RightAnkle abduction | 28.85 | 0.051 | 0.35 | 0.014 |
| RightAnkle rotation | 25.44 | 0.046 | 0.26 | 0.014 |
| LeftHip flexion | 13.87 | 0.064 | 0.43 | 0.010 |
| LeftHip abduction | 54.89 | 0.055 | 0.45 | 0.015 |
| LeftHip rotation | 44.03 | 0.098 | 0.69 | 0.015 |
| LeftKnee flexion | 22.77 | 0.080 | 0.44 | 0.016 |
| LeftKnee abduction | 47.87 | 0.043 | 0.41 | 0.012 |
| LeftKnee rotation | 58.09 | 0.061 | 0.46 | 0.016 |
| LeftAnkle flexion | 31.82 | 0.056 | 0.36 | 0.015 |
| LeftAnkle abduction | 7.29 | 0.081 | 0.24 | 0.015 |
| LeftAnkle rotation | 29.78 | 0.085 | 0.51 | 0.016 |
